# Linear regression of sampling distributions of the mean

**DOI:** 10.1101/2021.01.29.428767

**Authors:** David J Torres, Ana Vasilic, Jose Pacheco

## Abstract

We show that the simple and multiple linear regression coefficients and the coefficient of determination *R*^2^ computed from sampling distributions of the mean (with or without replacement) are equal to the regression coefficients and coefficient of determination computed with individual data. Moreover, the standard error of estimate is reduced by the square root of the group size for sampling distributions of the mean. The result has applications when formulating a distance measure between two genes in a hierarchical clustering algorithm. We show that the Pearson R coefficient can measure how differential expression in one gene correlates with differential expression in a second gene.

## Introduction

Linear regression coefficients and the Pearson R correlation has long been used to quantify the relationship between dependent and independent variables [1]. However, the “ecological fallacy” has shown that linear regression and correlation coefficients based on group averages cannot be used to estimate linear regression and correlation coefficients based on individual scores [2, 3].

It may not be well known that if all possible groups are considered, in the case of sampling distributions of the mean, the Pearson R coefficient computed from the group averages is equal to the Pearson R coefficient computed from the original individual scores for one independent variable [4, 5].

We extend this result and show that the linear regression coefficients (for simple and multiple regression) and the coefficient of determination *R*^2^ computed from sampling distributions of the mean (with or without replacement) are the same as the coefficient of determination and linear regression coefficients computed with the original individual data. The sampling distributions of the mean can also be constructed using differences between two groups of different size. The result has implications for hierarchical clustering of genes. Specifically, the Pearson R coefficient can be used to measure how differential expression in one gene correlates with differential expression in a second gene.

The standard error of estimate is a measure of accuracy for the surface of regression [6]. Using the coefficient of determination, we show that the standard error of estimate is reduced by the square root of the group size for sampling distributions of the mean.

In Section 1, we recall and reformulate the system of equations that are solved to determine the linear regression coefficients for individual scores. In Section 2, we prove the assertion that the same system of equations needs to be solved for sampling distributions of the mean with and without replacement or differences between two groups of sampling distributions. In Section 3, we show that the coefficient of determination is the same whether it is calculated using individual scores or all possible group averages from sampling distributions. Section 4 shows that the standard error of estimate is reduced by the square root of the group size for sampling distributions of the mean. Section 5 performs numerical simulations to illustrate these principles. Section 6 applies these results and shows that the Pearson R coefficient can be used to measure how differential expression in one gene correlates with differential expression in a second gene when the z-statistic is used.

## 1 Computing regression coefficients

Multiple regression requires one to compute the coefficients 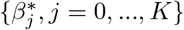 that minimize the sum of squares

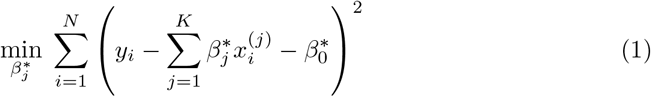

where *x*^(1)^, *x*^(2)^,…, *x*^(*K*)^ represent *K* different independent variables, *y* is the dependent variable, and the *i^th^* realization of variables *x*^(*j*)^ and *y* are 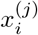 and *y_i_*, respectively, 1 ≤ *i* ≤ *N*. Note that for simple linear regression, *K* = 1. We recast the sum (1) in the form

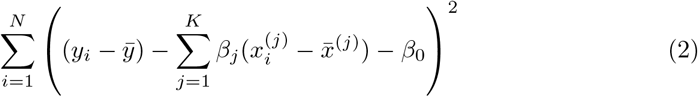

where

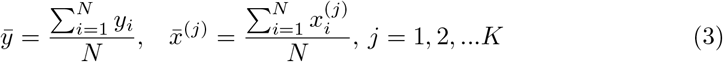

and the coefficients are related by

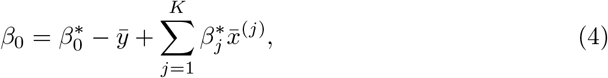

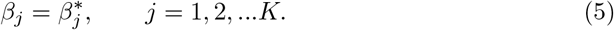

To solve for *β*_0_, we set the partial derivative of (2) with respect to *β*_0_ to zero which yields

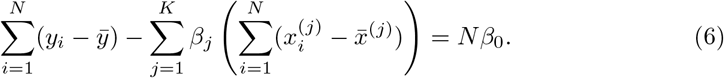

However

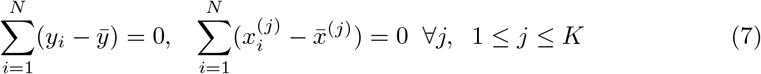

which implies that *β*_0_ = 0. Thus one can redefine the problem of computing multiple regression coefficients (1) to be selection of the coefficients {*β_j_, j* = 1,…, *K*} that minimizes the sum of squares

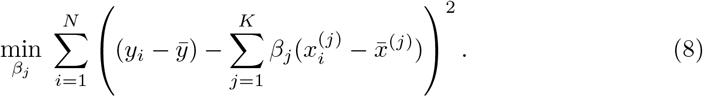

In the matrix approach to minimizing the sum of squares which can be derived by setting the partial derivatives 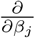 of (8) to zero, the system of equations in (8) is written in matrix form [7]

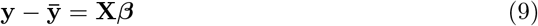

where

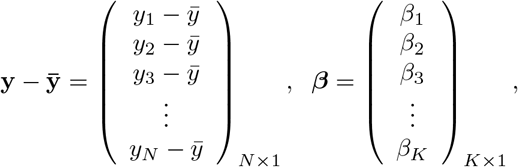

and **X** is a *N* by *K* matrix whose entries are:

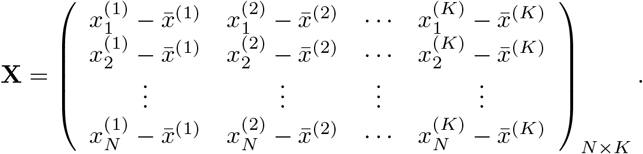

One can solve for the multiple regression coefficients in the vector ***β*** by left multiplying (9) by the transpose **X**^*T*^ and solving the linear system

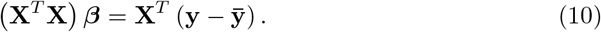

The elements in the square *K* by *K* matrix **X**^*T*^**X** will be sums of the form

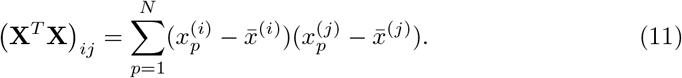

Similarly the entries in the *K* by 1 vector 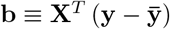 will be sums of the form

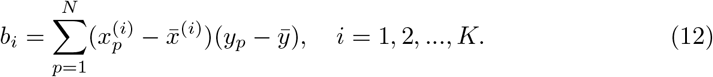

It should be noted that for each pair of fixed indices *i* and *j*, the sum in either expression (11) or (12) can be represented using a sum of the form

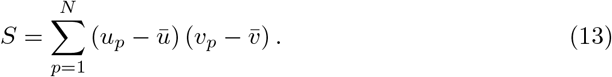

In the following section, we show that if the variables *x*^(1)^, *x*^(2)^,…, *x*^(*k*)^, and *y* are replaced with all the elements from the sampling distributions of the mean, the system (14) is obtained

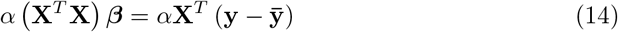

for some constant *α*. Moreover, we obtain a closed form for the constant *α*. If *m* is the group size and we account for order, the size of the matrix **X** will be *N^m^* × *K* for selections with replacement and 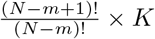 for selections without replacement. However, the resulting system (14) will still be a *K* × *K* system. Since the system (14) is equivalent to the system (10), the regression coefficients for the sampling distributions of the mean will be the same as the regression coefficients computed from the original data according to (5) for 1 ≤ *j* ≤ *K*. The equivalence of 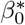 follows from *β*_0_ = 0, equation (4), and the fact that the means of the original data (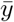 and 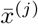) are the same as the means computed using all the elements from the sampling distributions of the mean (with or without replacement). If we assume that there are *N_p_* elements in the sampling distribution, this can be stated mathematically as

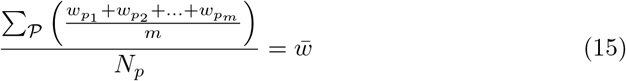

or

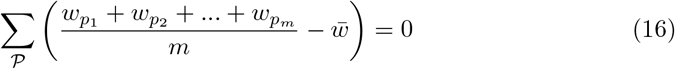

where *w* can represent *y* or *x*^(*j*)^ and 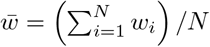. The sum 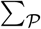 is a sum over all possible index values in the sampling distribution.

## 2 Regression with averages

Let us create elements from the sampling distribution of the mean using elements chosen from the groups

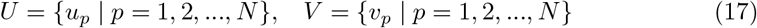

by averaging all possible groups of size *m*_1_ and size *m*_2_

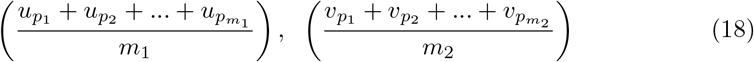

chosen from the sets *U* and *V*. We assume without loss of generality that *m*_1_ ≥ *m*_2_. The first *m*_2_ choices are paired

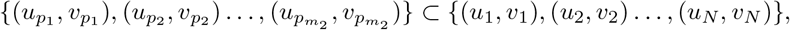

while the remaining choices remain unpaired

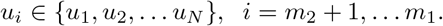

If the selections are done without replacement, *p_r_* ≠ *p_s_* if *r* ≠ *s*. However, if the selections are formed with replacement, *p_r_* can equal *p_s_*.

Let us now replace *u_p_* and *v_p_* in (13) with all possible averages of *m*_1_ and *m*_2_ elements as shown in (18),

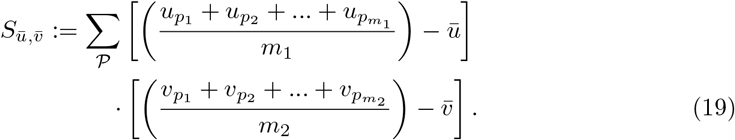

where

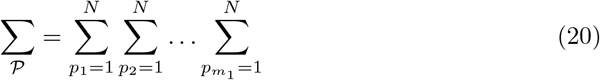

for selections with replacement and

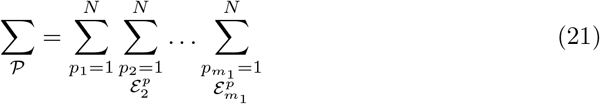

for selections without replacement where 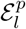 is used to denote the exclusion of previously chosen indices

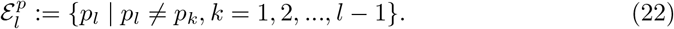

The means of the original scores 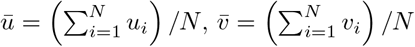 are used in (19) since they are equal to the means of the sampling distributions by (15). Note that order matters in the way the sums are written in (20-of the mean share the same regression plane 21). For example, (*u*_1_, *u*_2_, *u*_3_) is considered a different choice then (*u*_3_, *u*_2_, *u*_1_). Disregarding order would lead to *m*_1_! fewer terms in (21). However the same system (14) would be generated if order was not considered for sampling distributions without replacement.

Factoring out 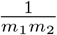 from (19) yields

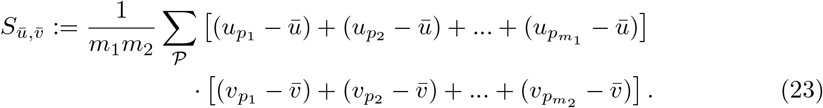

Sections 2.1 and 2.2 show that 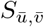 will be equal to a factor *α* times *S* as defined by (13) for sampling distributions with and without replacement respectively. Section 2.3 generalizes these results to differences of two groups of sampling distributions. In all cases, the elements of the matrix (**X**^*T*^**X**) and the vector 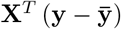 will be multiplied by the same factor *α* when elements from the sampling distributions of the mean are used.

### 2.1 Sampling distributions with replacement

Start with 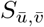 as defined by (23). Since we are considering sampling distribution with replacement, the values chosen for summation indices *p_i_* do not need to be different. We will show that

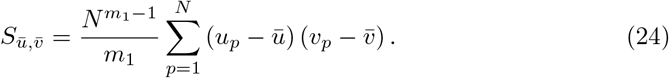

If we distribute the sums inside the parentheses in (23), two types of terms are formed. The first type of term takes the form

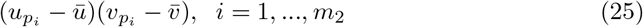

where the same summation index *p_i_* is used for 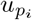 and 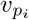. The second type of term takes the form

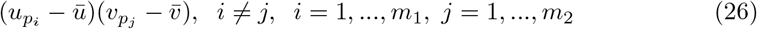

where different summation indices, *p_i_* and *p_j_* are used.

All the terms of the form shown in (26) are zero since when the sums 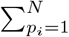 and 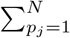 from 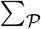 are moved to apply directly to 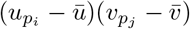, each term can be summed independently

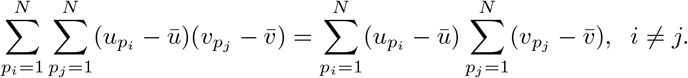

However

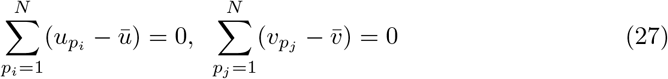

as noted by equation (7). Thus we must only consider terms of the form (25). The sum (20) acting on 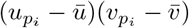 can be rearranged as

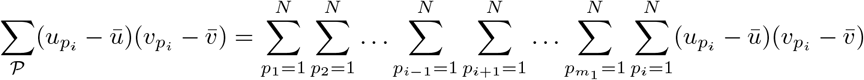

and simplified to

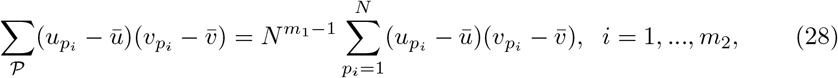

since each sum 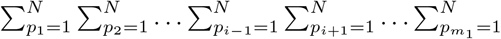 contributes a factor of *N*. Multiplying the right side of (28) by *m*_2_ to ensure all summation indices *p_i_, i* = 1,…, *m*_2_ are accounted for and multiplying by the factor 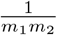 present in (23) yields (24). One can also derive (24) using random variables and expected values.

Since 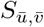 is a multiple of *S* defined by (13), the system of equations (14) will be formed where 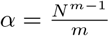 when we set *m* = *m*_1_ = *m*_2_. Thus the multiple regression coefficients ***β*** computed from sampling distributions of the mean with replacement will be equal to the multiple regression coefficients computed from the original scores.

### 2.2 Sampling distribution without replacement

We will show that

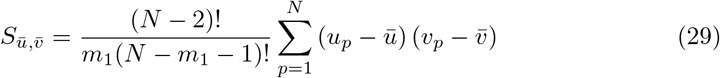

for sampling distributions created without replacement. If we distribute the sums inside the parentheses of (23), we again distinguish between two types of terms: terms of the form 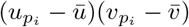, *i* = 1,…, *m*_2_ and terms of the form 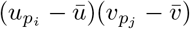, *i* ≠ *j, i* = 1,…, *m*_1_, *j* = 1,…, *m*_2_.

Choose a summation index *p_i_*. The sum (21) applied to 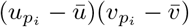 can be written as

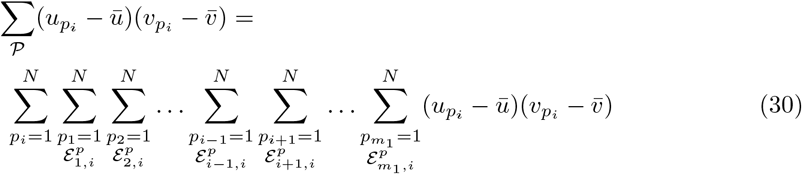

where the sum 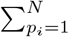 with summation index *p_i_* is placed first and the term

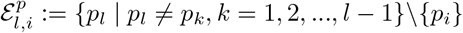

excludes previously chosen index values and the index value chosen for *p_i_*. The right side of (30) can be simplified to

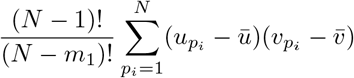

since the choice made for *p_i_* in the first sum 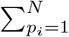 leaves *N* − 1 choices for the second sum, *N* − 2 choices for the third sum, up to *N* − (*m*_1_ − 1) choices for the last sum.Moreover there are *m*_2_ terms similar to (30) for each summation index, *p_i_*. Thus the terms of the form 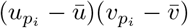 contribute

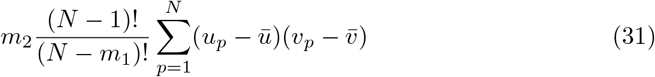

to the sum 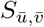.

We now consider terms of the form 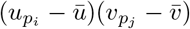 with two different summation indices *p_i_* and *p_j_*. The sum (21) applied to 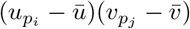, *i* ≠ *j* can be written as

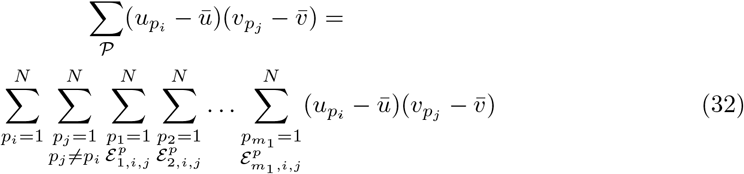

where the sums 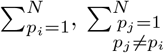 with summation indices *p_i_* and *p_j_* are placed first and the term

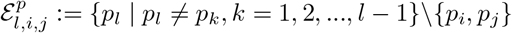

excludes previously chosen index values and the index values chosen for *p_i_* and *p_j_*. Equation (32) can be simplified to

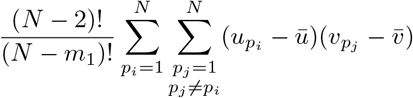

since the choice made for *p_i_* in the first sum 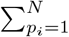 and *p_j_* in the second sum 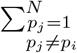 leaves *N* − 2 choices for the third sum, *N* − 3 choices for the fourth sum, up to *N* − (*m*_1_ − 1) choices for the last sum. Moreover there are *m*_2_(*m*_1_ − 1) sums of the form (32) that can be identified when the terms in (23) are distributed. Therefore the terms of the form 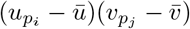 contribute

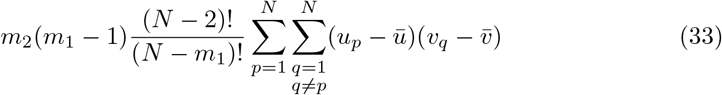

to the sum 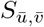. Remove 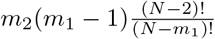 terms of the form 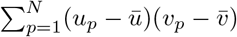 from (31) and add them to (33) to form

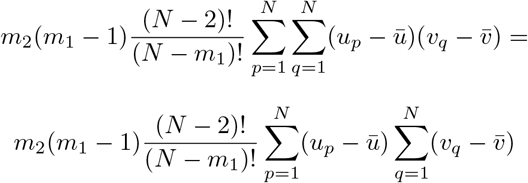

which is zero by (27). This leaves

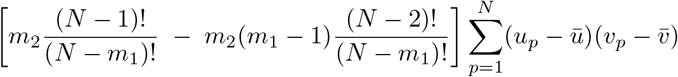

remaining terms from (31) which simplifies to (29) after multiplying by the factor 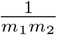 present in (23)

Since 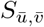 is a multiple of *S* as defined by (13), the system of equations (14) will be formed where 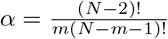 when we set *m* = *m*_1_ = *m*_2_. Thus the multiple regression coefficients ***β*** computed from sampling distributions of the mean without replacement will be equal to the multiple regression coefficients computed from the original scores.

### 2.3 Difference between two groups

The results in Sections 2.1 and 2.2 generalize to a difference of two groups of sampling distributions. Let *m*_1_ be the size of Group 1 and *m*_2_ be the size of Group 2. The two groups can be composed to allow or exclude common elements.

#### 2.3.1 Group 1 and Group 2 can share elements

Consider the expression *S_d_* shown in (34). The sum 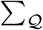 composed of *m*_2_ iterated sums is essentially the same sum shown in either (20) or (21) except that the indexing is done with *q* instead of *p*. We first examine the case where Group 1 and Group 2 can share elements: i.e. *p_i_* may be equal to *q_j_*.

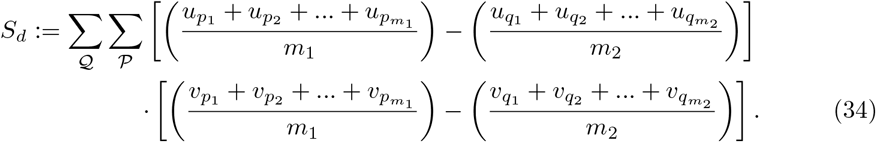

*S_d_* can be written in the form

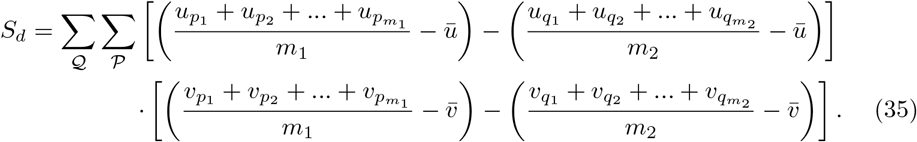

Distributing gives

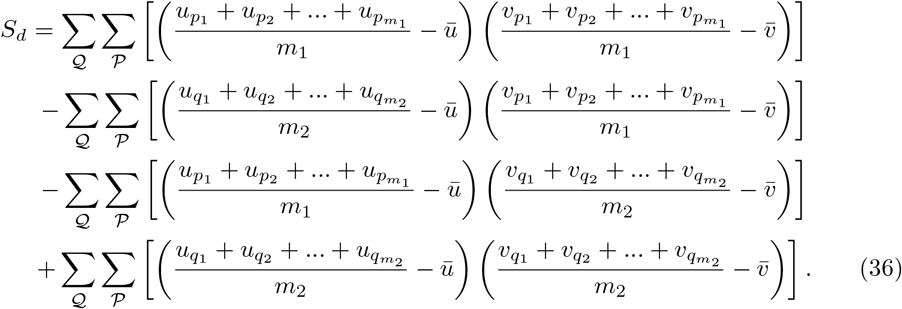

After one accounts for the sums 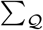 in the first term and 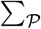 in the fourth term, one can write

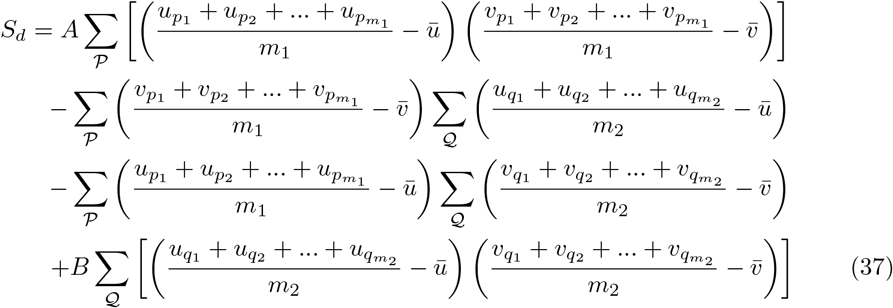

where

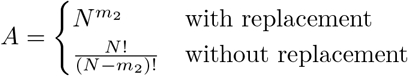

and

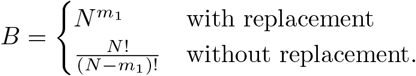

The expression for *A* can be derived by recognizing that *N* choices are available for each of the *m*_2_ sums in 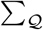 when the selections are made with replacement. When the selections are made without replacement, there are *N* choices for the first sum, *N* − 1 choices for the second sum, up to *N* − (*m*_2_ − 1) for the *m*_2_’th sum. The same reasoning can be used to derive the expression for *B*. By (16), the second and third terms in (37) are zero and can be eliminated. Using equations (24) and (29), one can simplify (37) to

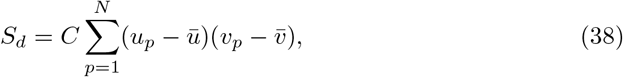

where

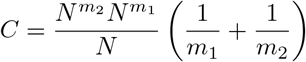

when selections are made with replacement and

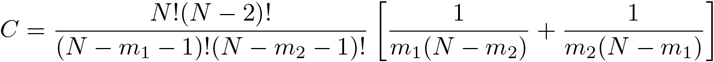

when selections are made without replacement.

Note that the mean of a difference of two groups of sampling distributions of the mean is zero. When 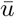 and 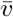 are set to zero in (19) and a difference of two groups of sampling distributions are used, it is evident that *S_d_* is similar in format to (19). Thus the system of equations (14) will be formed where *α* = *C*. Thus the multiple regression coefficients ***β*** computed from a difference of two groups of sampling distributions of the mean will be equal to the multiple regression coefficients computed from the original scores.

#### 2.3.2 Group 1 and Group 2 do not simultaneously share elements

We also consider the case where Group 1 and Group 2 do not simultaneously share any elements. We assume the selections are done without replacement. Under these restrictions, one can write (36) as

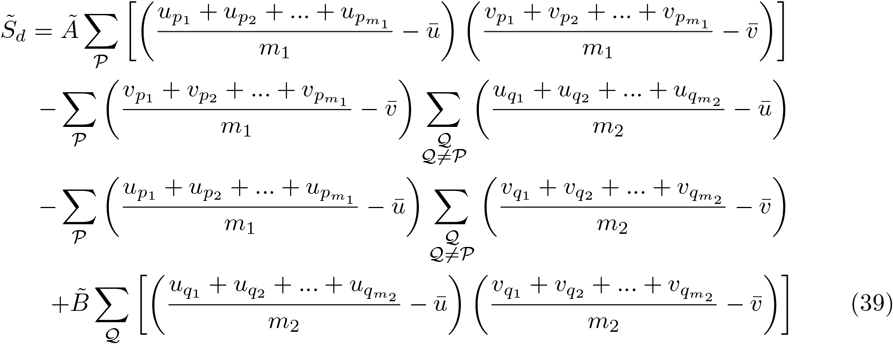

where

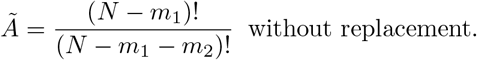

and

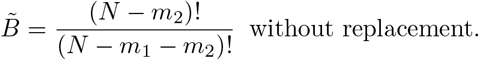

The notation 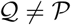 is used to exclude any elements in the sum 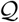 from indices previously selected in the sum 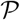. The *Ã* coefficient can be derived by noting that for the distinct *m*_1_ indices 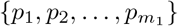 chosen in 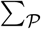, there remain *N* − *m*_1_ choices for the first sum in 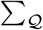, *N* − *m*_1_ − 1 choices for the second sum, and so on up to *N* − *m*_1_ − (*m*_2_ − 1) choices for the *m*_2_’th sum. Similarly, the 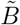 coefficient can be derived by noting that for the distinct *m*_2_ indices 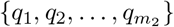 chosen in 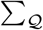, there remain *N* − *m*_2_ choices for the first sum in 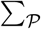, *N* − *m*_2_ − 1 choices for the second sum, and so on up to *N* − *m*_2_ − (*m*_1_ − 1) choices for the *m*_1_’th sum. Turning to the second half of the second term of (39), which we define to be

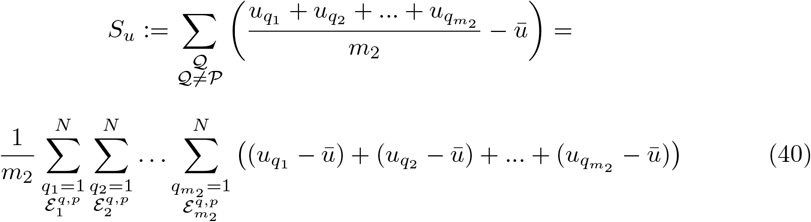

where

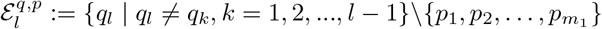

excludes previously chosen indices in the 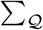 sum and any previously chosen indices 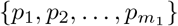 selected from the 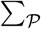 sum. Applying the sum to the specific term 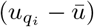 the sum 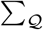 can be rearranged as

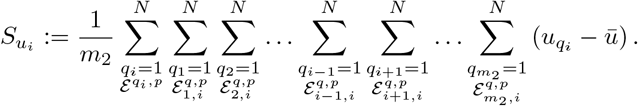

where the term

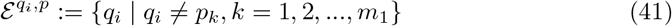

excludes previously chosen indices 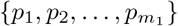 selected from the sum 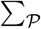 and where the term

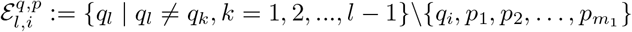

excludes previously chosen indices in the 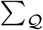 sum, the index *q_i_* chosen in the 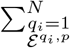 sum, and any previously chosen indices 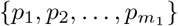 selected from the 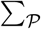 sum. Bear in mind that the sums

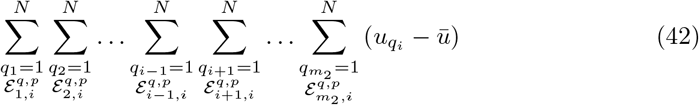

will contribute the same factor to 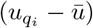 regardless of the selected value for the summation index *q_i_*. Taking care to avoid selecting a index that has been already chosen, we note that *m*_1_ choices have already been made for the set 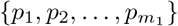. In addition, for each choice of *q_i_* in 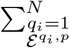 there remain *N* − *m*_1_ − 1 choices left for the first sum 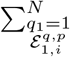, *N* − *m*_1_ − 2 choices left for the second sum, up to *N* − *m*_1_ − (*m*_2_ − 1) choices for the (*m*_2_ − 1)’th sum or

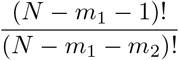

total choices. Thus

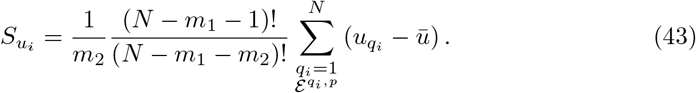

Using the definition of the excluded terms 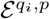 (41) in the sum 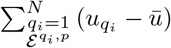,

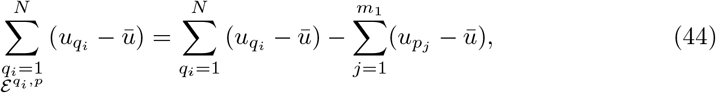

one can replace 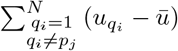 in (43) with the right hand side of (44) to yield,

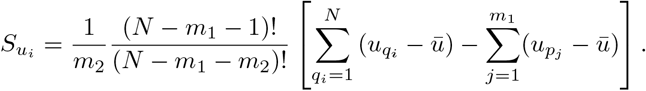

The sum 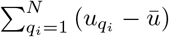 is zero by (27). Since there are *m*_2_ terms of the form 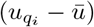 in (40), *S_u_* can be written as

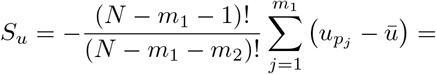

or

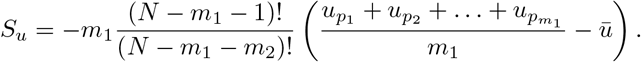

We can apply the same steps to the second half of the third term of (39), which we define to be *S_v_*

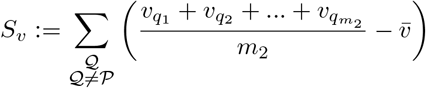

to show that

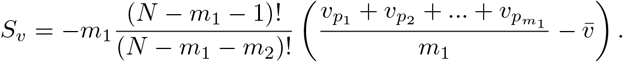

Using these results in (39),

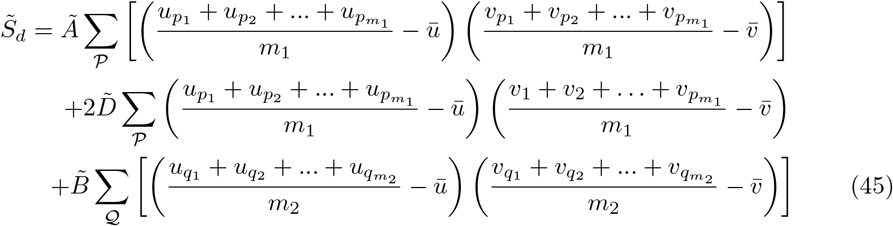

where

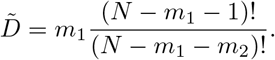

Using equation (29), (45) simplifies to

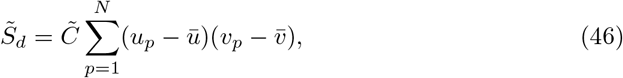

where

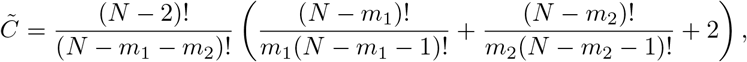

keeping in mind that the selections are made without replacement. Again 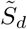 is a multiple of *S*. Therefore the system of equations (14) will be formed where 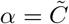.

## 3 Coefficient of determination

The coefficient of determination *R*^2^ is the proportion of variability in the dependent variable that can be accounted for by the independent variables [6]. It is defined using

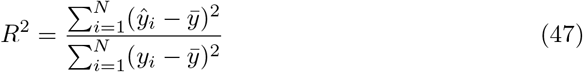

where *ŷ_i_* is the prediction provided by the surface of regression

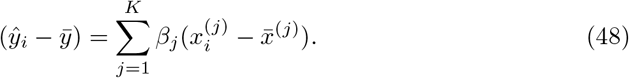

Substituting (48) into (47),

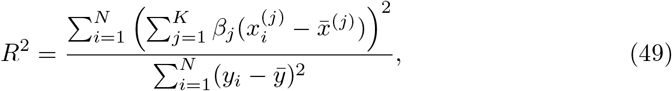

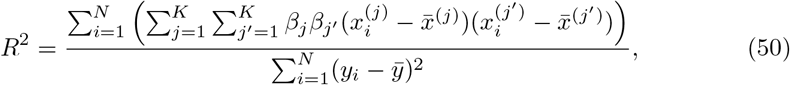

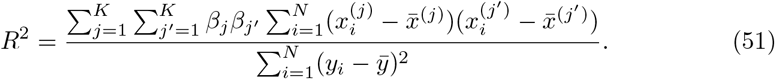

Again we see the presence of sums 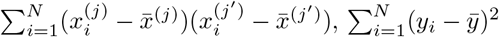 of the form (13). Both numerator and denominator will be multiplied by the same constant according to (24), (29), (38), and (46) leaving the coefficient of determination *R*^2^ unchanged when elements of the sampling distribution of the mean or differences of two groups of sampling distributions of the mean are used.

## 4 Standard error of estimate

The standard error of estimate is a measure of accuracy for the surface of regression [8]. In this section, we show the standard error of estimate is reduced by the factor 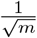 where *m* is the group size for sampling distributions with replacement and by

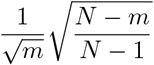

for sampling distributions without replacement.

The sum of squares error *SSE* is defined to be

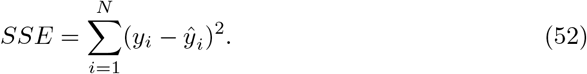

Given this definition, the standard error of estimate *s_e_* can be defined

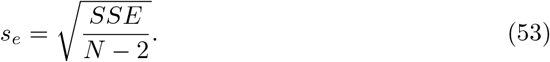

Now by [7]

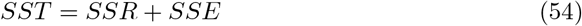

where SST is the total variation and SSR is the sum of squared regression,

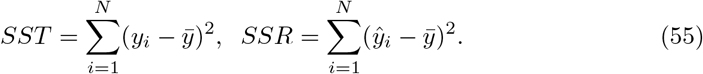

With these definitions, the coefficient of determination (47) can also be written as

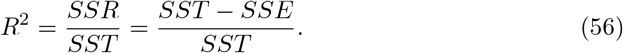

Solving (56) for SSE and dividing by *N*

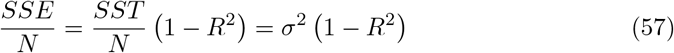

where

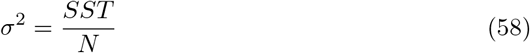

is the population variance. Now by (53) 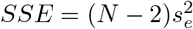. Replacing SSE in (57) with 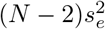 and solving for *s_e_* yields

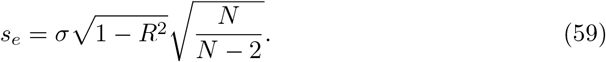

When sampling distributions of the mean are used, *R* remains the same, but *σ* is replaced by 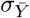 where

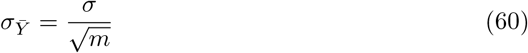

for sampling distributions with replacement and

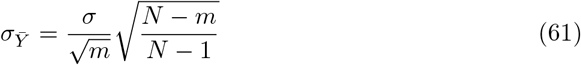

for sampling distributions without replacement [8]. Thus for selections made with replacement and for selections made without replacement (if *N* >> *m*), *s_e_* will be reduced by 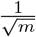 when sampling distributions of the mean are used. This result is analogous to the reduction of the standard deviation by 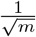 when using sampling distributions of the mean for one variable.

## 5 Numerical simulations

Figure 1 plots the original data {(*x_i_, y_i_*), 1 ≤ *i* ≤ 10} in red and all elements from the sampling distribution of the mean generated without replacement for groups of size *m* = 2 in blue and *m* = 3 in green for *N* = 10 original points. The original data and elements from the sampling distribution of the mean share the same regression line and coefficient of determination *R*^2^. The elements of the sampling distribution of the mean are clustered more closely about the regression line compared to the original data which is consistent with (59) and (61).

**Fig 1.**
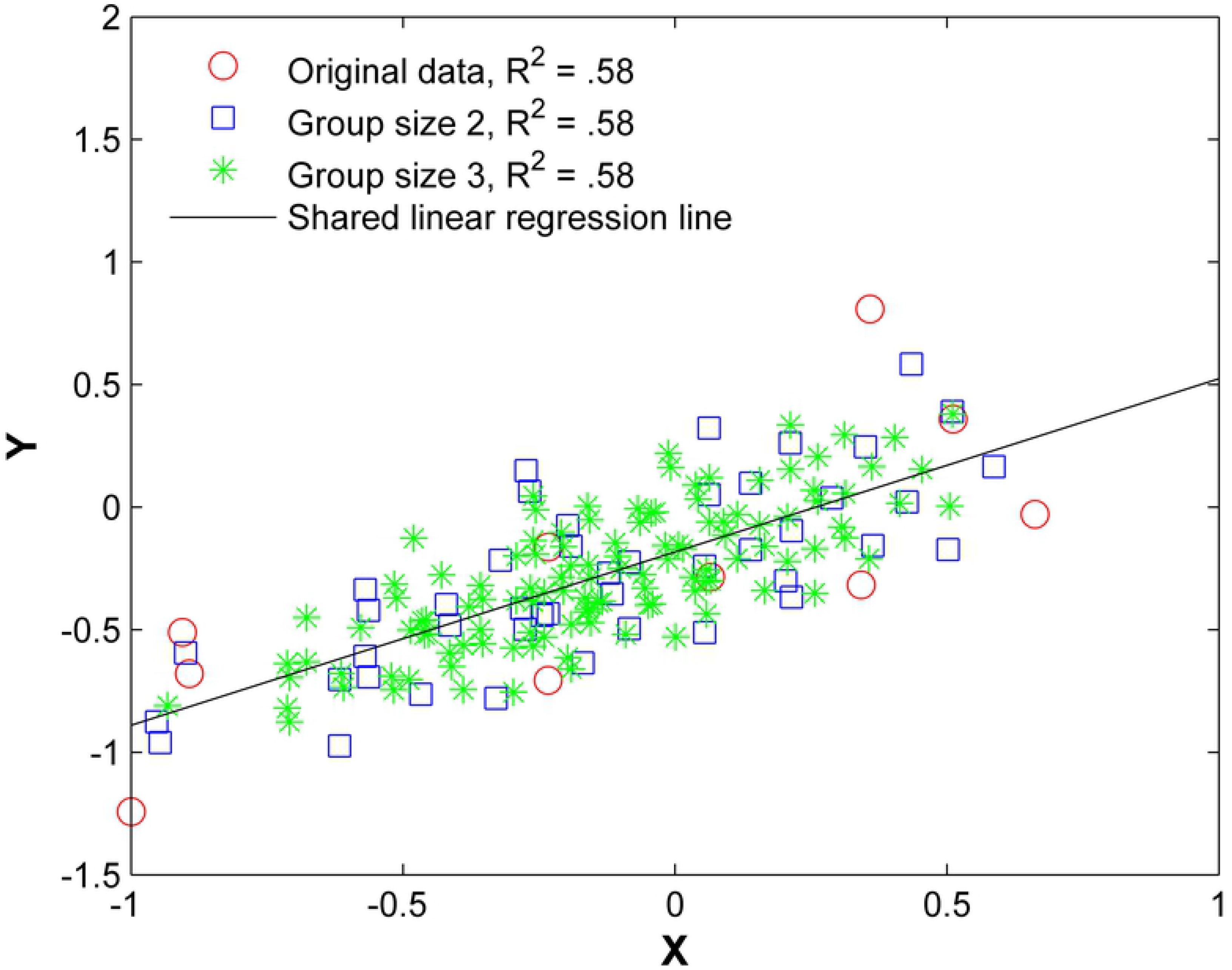
Original data (*x_y_, y_i_*), all elements from the sampling distributions of the mean, and the shared linear regression line. The red circles are the original 15 points, the blue squares are the averaged data of size *m* = 2, and the green asterisks are the averaged data of size *m* = 3 without replacement.

Figure 2 plots the original data (*x_i_, y_i_, z_i_*), 1 ≤ *i* ≤ 15 in red and all elements from the sampling distribution of the mean generated without replacement for groups of size *m* = 2 in blue and *m* = 3 in green for *N* = 15 original points. The original data and elements from the sampling distribution of the mean share the same regression plane *z* = .653*x* − .712*y* and coefficient of determination *R*^2^ = 0.76. For visualization purposes, the normal distance to the plane is plotted as the z-coordinate and the multiple regression plane is aligned with the *z* = 0 plane.

**Fig 2.**
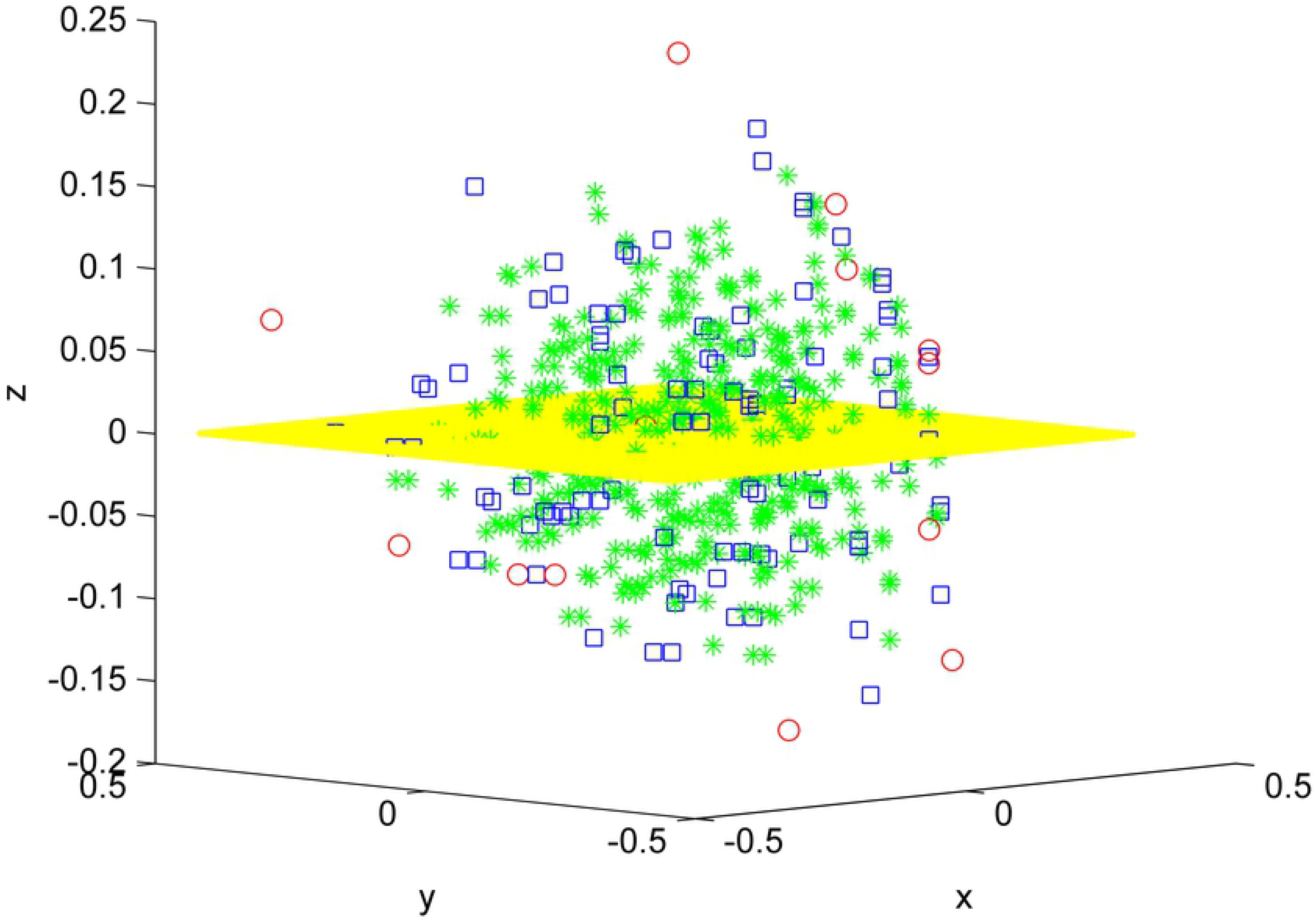
Original data (*x_i_, y_i_, z_i_*), *i* = 1, 15 in red and all elements from the sampling distribution of the mean for *m* = 2 (blue) and *m* = 3 (green), and the shared linear regression plane.

Figure 3 shows the convergence of sampling distributions of the mean for {(*x_i_, y_i_*), *i* = 1, 2,…, *N*}, *N* = 11 scores with Pearson correlation coefficient *R* = 0.35 and linear regression slope *β*_1_ = 0.27. In the first simulation shown in black and red, elements from the sampling distributions of the mean are created using groups of size *m* = 5 without replacement. In the second simulation shown in blue and green, elements from the sampling distributions of the mean are created using differences of two groups of size *m*_1_ = 4 and *m*_2_ = 2 without replacement. The horizontal axis plots the fraction of total selections used in the sampling distributions. There are 11^5^ = 161, 051 total selections for the first simulation and 11!/5! = 332, 640 total selections for the second simulation. The vertical axis plots the base 10 logarithm of the absolute difference. The absolute difference can be between either the Pearson R = 0.35 based on individual scores and the Pearson R computed from a fraction of the elements from the sampling distributions of the mean (black and blue graphs), **or** between the linear regression slope *β*_1_ = .27 based on the original scores and the slope computed using from a fraction of the elements from the sampling distributions of the mean (red and green graphs). While not entirely obvious due to the density of points, all differences decrease from approximately 10^−6^ to less than 10^−13^ in the last 0.001% of the total selections. In addition, the differences do not always decrease monotonically as the fraction of total selections increase, and the differences decrease to very small values (less than 10^−5^) at certain points during the course of the convergence as noted by the downward spikes.

**Fig 3.**
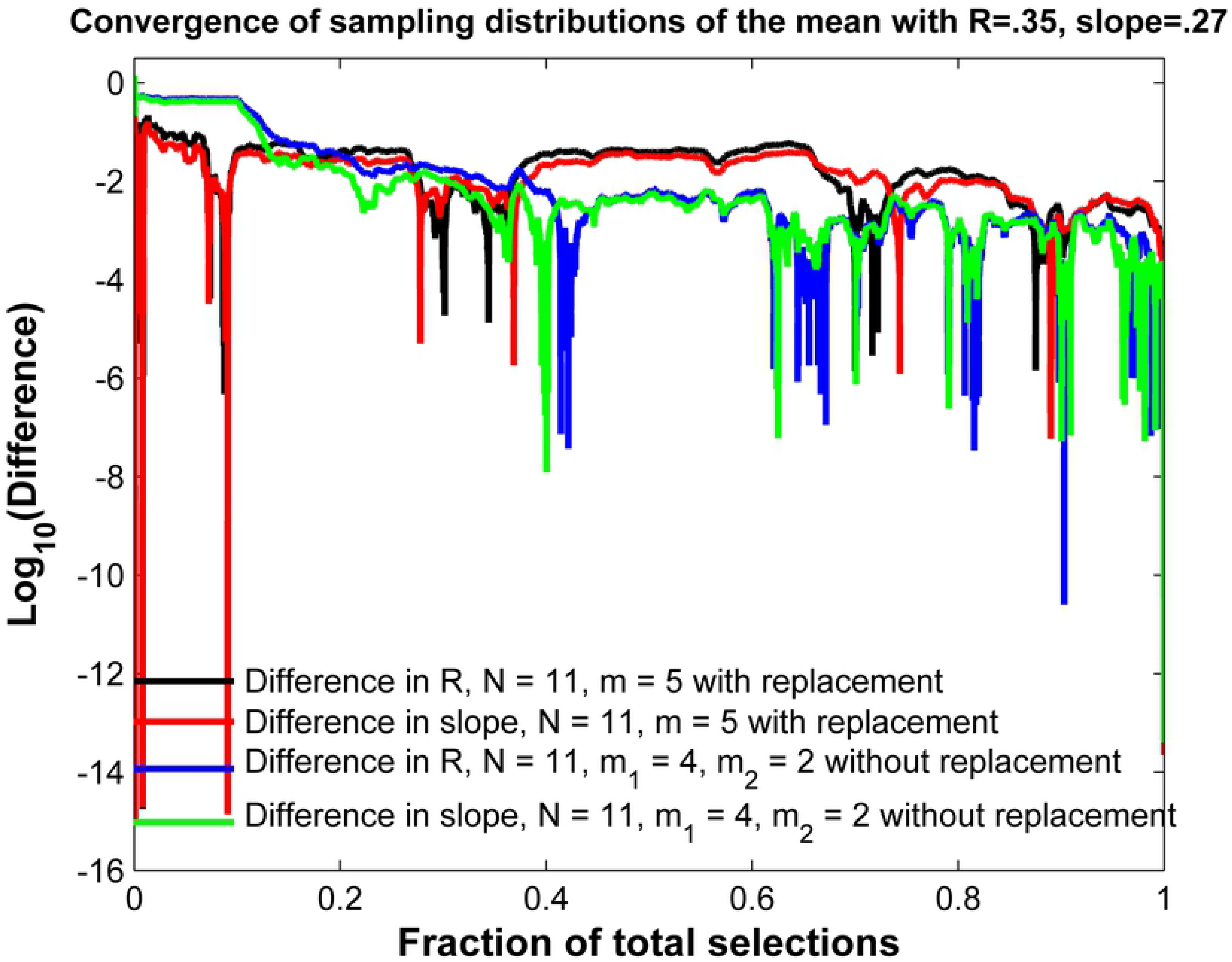
Convergence of the Pearson *R* and linear regression slope for sampling distributions of the mean.

## 6 Gene expression and distance between genes

A useful way of organizing the data obtained from microarrays or RNA-seq data is to group together genes that exhibit similar expression patterns through hierarchical clustering. A hierarchical clustering algorithm generates a dendrogram (tree diagram). However, the algorithm requires that a distance be defined to quantify similarities in expression between two individual genes.

Let *A_i_* denote the expression level of gene *A* for patient *i* and let *B_i_* denote the expression level for gene *B* for patient *i*, 1 ≤ *i* ≤ *N*. Distances between genes can be computed using many metrics [9], but two common ones are the Euclidean distance

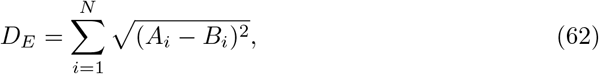

and the Manhattan distance,

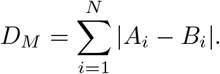

Correlation coefficients [10] are also used to measure the similarities between two genes. One measure of distance using the Pearson R is

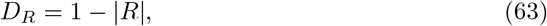

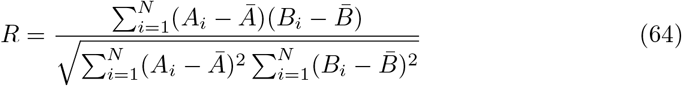

or 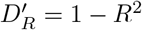 [11] if the sign of *R* is not important. If *R* is close to 1 or −1, the distances *D_R_*, 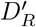 will be close to zero.

The purpose of the next section is to propose a new distance based on the differential expression of two genes. We then show the new measure of distance is the same as the Pearson R coefficient computed from the original scores (64), thus lending support to the use of the Pearson R coefficient in measuring the distance between two genes.

### 6.1 Formulating a new distance between two genes

Let us formulate a new distance based on differential expression. Select 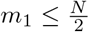 distinct random patients and their expression levels for gene *A* and assign them to Group 1. Select a second group of 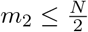 distinct (and different from Group 1) random patients and assign their expression levels to Group 2. Repeat the process using the same selections for gene *B*. Since both groups are sampled from a population with a known variance *σ*^2^, the z-statistic [12] for two independent samples can be used to measure differential expression for gene A

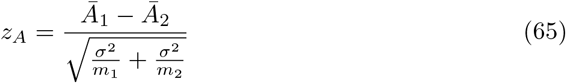

which if *m*_1_, *m*_2_ ≥ 30 will be approximately normally distributed. Let *z_B_* be the z-statistic for gene *B* for the same selection of patients using the same equation (65). This process can be repeated multiple times giving a set of ordered pairs 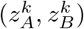 for each different selection (*k*) of groups. The Pearson *R* value, *R_t_* can then be computed from these ordered pairs using all possible selections *K*

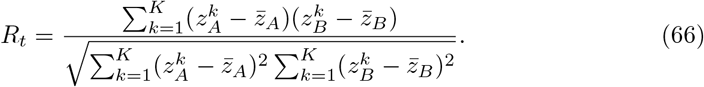

The new distance will now be defined as *D_T_* or alternatively 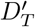

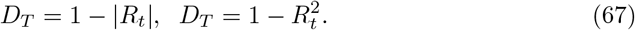

Given *N* total patients, there exist 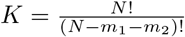 total selections. Computing all selections is prohibitive for large *N*. However, we know from the analysis in Section 2.3.2, and the fact that the Pearson *R* coefficient is not affected by the multiplicative factor 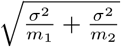 in (65), that the distance *D_R_* will be equal *D_T_* and 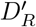 will be equal 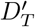.

## 7 Conclusion

We have shown that the linear regression coefficients (simple and multiple) and the coefficient of determination *R*^2^ computed from sampling distributions of the mean (with or without replacement) are equal to the regression coefficients and coefficient of determination computed with the original data. This result also applies to a difference of two groups of sampling distributions of the mean. Moreover, the standard error of estimate is reduced by the square root of the group size for sampling distributions of the mean.

The result has implications for the construction of hierarchical clustering trees or heat maps which visualize the relationship between many genes. These processes require one to define a distance between two genes using their expression levels. We developed a new measure of distance based on how differential expression in one gene correlates with differential expression in a second gene using the z-statistic. We showed that the new measure is equivalent to the Pearson R coefficient computed from the original scores, thus lending support to the use of the Pearson R coefficient for measuring a distance between two genes.

## References

1. Yan X. Linear regression analysis: Theory and computing. Singapore: World Scientific; 2009.

2. Robinson WS. Ecological correlations and the behavior of individuals. Am Sociol Rev. 1950; 15(3):351–357, https://doi.org/10.2307/2087176.

3. Goodman LA. Ecological regressions and the behavior of individuals. Am Sociol Rev. 1953; 18:663–664.

4. Kenney JF. Mathematics of statistics, Part Two. New York: D. Van Nostrand Company, Inc.; 1947.

5. Pearson K. On the probable errors of frequency constants. Biometrika. 1913; 9:1–10.

6. Pagano R. Understanding statistics in the behavioral sciences, 10th ed. Belmont, California: Wadsworth Publishing; 2012.

7. Walpole RE and Myers RH. Probability and statistics for engineers and scientists, 3rd ed. New York: MacMillan Publishing Company; 1985.

8. Sullivan M III. Fundamentals of statistics, 3rd ed. New York: Pearson; 2010.

9. D’haeseleer P. How does gene expression clustering work? Nat Biotechnol. 2005; 23:1499–1501.

10. Quackenbush J. Computational analysis of microarray data. Nat Rev Genet. 2001; 2(6):418–427.

11. Eisen MB, Spellman PT, Brown PO, Botstein D. Cluster analysis and display of genome-wide expression patterns. Proc Natl Acad Sci USA. 95(25):14863–14868.

12. Triola MF. Elementary statistics using the TI-83/84 Plus calculator, 3rd ed. New York: Addison-Wesley; 2011.

